# Estimating gene conversion rates from population data using multi-individual identity by descent

**DOI:** 10.1101/2025.02.22.639693

**Authors:** Sharon R. Browning, Brian L. Browning

## Abstract

In humans, homologous gene conversions occur at a higher rate than crossovers, however gene conversion tracts are small and often unobservable. As a result, estimating gene conversion rates is more difficult than estimating crossover rates. We present a method for multi-individual identity-by-descent (IBD) inference that allows for mismatches due to genotype error and gene conversion. We use the inferred IBD to detect alleles that have changed due to gene conversion in the recent past. We analyze data from the TOPMed and UK Biobank studies to estimate autosome-wide maps of gene conversion rates. For 10 kb, 100kb, and 1 Mb windows, the correlation between our TOPMed gene conversion map and the deCODE sex-averaged crossover map ranges from 0.56 to 0.67. We find that the strongest gene conversion hotspots typically die back to the baseline gene conversion rate within 1 kb. In 100 kb and 1 Mb windows, our estimated gene conversion map has higher correlation than the deCODE sex-averaged crossover map with PRDM9 binding enrichment (0.34 vs 0.29 for 100 kb windows and 0.52 vs 0.34 for 1 Mb windows), suggesting that the effect of PRDM9 is greater on gene conversion than on crossover recombination. Our TOPMed gene conversion maps are constructed from 55-fold more observed allele conversions than the recently published deCODE gene conversion maps. Our map provides sex-averaged estimates for 10 kb, 100 kb, and 1 Mb windows, whereas the deCODE gene conversion maps provide sex-specific estimates for 3 Mb windows.

## Introduction

In meiosis, the two haplotypes of a parent are combined via recombination to produce the gamete’s haplotype. Recombination takes the form of crossovers and gene conversions. Crossovers are positions at which switches occur between the two haplotypes, and the average distance between crossovers is approximately 100 million base pairs in humans.^1^ In homologous gene conversion, a tract of tens or hundreds of base pairs is copied onto the transmitted haplotype from the parent’s other haplotype.^2^ Gene conversion changes the allele on the transmitted haplotype only at positions where alleles on a parent’s two haplotypes differ, that is, at positions of heterozygosity in the parent. Since the heterozygosity rate in human populations is around 1 per thousand base pairs,^3^ many gene conversions are not observable. If an allele on the transmitted haplotype is changed by gene conversion, it can be difficult to determine whether the changed allele is due to gene conversion or genotype error.^4^

One approach to studying gene conversions is sperm-typing.^2^ By typing many sperm from one or more fathers, one can determine the haplotype phase of the fathers and detect possible alleles changed by gene conversion in the gametes. An advantage of this approach is that many meioses can be observed. A disadvantage is that genotype errors can produce miscalled alleles that look like alleles changed by gene conversion.^4^ In contrast, the use of multi-generational families enables resolution of genotype error as well as phase determination.^4; 5^ The use of nuclear families with more than one sibling does not address genotype error but does allow for phase determination.^6^ Collecting a large number of families is challenging. The largest such analysis to date included 10,840 meioses from 2132 nuclear families and identified 62,762 alleles changed by gene conversion. The resulting gene conversion maps provide estimated gene conversion rates in 3 Mb windows.^6^

Population-genetic models such as the coalescent can be used to investigate rates of gene conversion from genetic data from unrelated individuals.^7; 8^ The resolution of such methods is not high; for example, the resolution of one such analysis is one estimated rate per chromosome.^9^

In 2024, we proposed the use of multi-individual identity by descent (IBD) to detect alleles changed by gene conversion in population data.^10^ Application to a data from 125,361 individuals found 9,313,066 alleles changed by gene conversion, which was 2877 times as many allele conversions compared to the largest family study at that time, and 148 times as many allele conversions compared to a recently published study, which is the largest family study to date.^5; 6^ Our approach to detecting alleles changed by gene conversion from IBD data is robust to genotype error, because it requires that each allele conversion be observed in two or more identical-by-descent individuals. However, the IBD segment detection in our earlier method does not account for discordant alleles caused by genotype error or gene conversion. As a result, the power to detect IBD segments in regions with a high rate of gene conversion is reduced, which reduces power to detect allele conversions in these regions. This limitation makes our earlier method unsuitable for estimating gene conversion rates. Although our earlier approach can detect alleles changed by gene conversion, the analysis is cumbersome. Disjoint sets of markers must be used for detecting IBD and for detecting allele conversion, and multiple analyses with different marker sets must be performed in order to interrogate all markers for allele conversions.

In this work, we present a new method for detecting IBD that is robust to discordant alleles and that eliminates the need for multiple analyses with different sets of markers. Our new method, implemented in the ibd-cluster software package, employs a probabilistic model that accounts for genotype error and other sources of discordant alleles. We apply the method to infer 17,404,902 alleles changed by gene conversion across two data sets of sizes 39,961 and 125,361 individuals. We use the detected allele conversion to estimate the gene conversion rate at resolutions of 10 kb, 100 kb, and 1 Mb. We estimate the probability that a position in the genome is part of a gene conversion tract, rather than the gene conversion tract initiation rate. If the average length of gene conversion tracts is the same throughout the genome, these two rates will be proportional to each other. Our method estimates the relative rate of gene conversion as the rate varies along the genome; it does not estimate the genome-wide rate, which can be obtained from other pedigree-based or IBD-based methods.^4; 11^

## Subjects and Methods

### Multi-individual IBD inference

A set of haplotypes form an IBD cluster at a locus if they share a recent common ancestor. In inferring multi-individual IBD, i.e. IBD clusters, there is an implicit or explicit dependence on underlying pairwise IBD segments.^10; 12; 13^ However, in spite of this dependence, the inferred multi-individual IBD is not easily described in terms of segments and is more readily expressed as sets of IBD haplotypes at a locus.^10^ Locus-based multi-individual IBD is ideal for detecting alleles changed by gene conversion.^10^

We previously developed a method for multi-individual IBD inference that can be applied to large samples of individuals.^10^ The method did not allow for discordant alleles in IBD sequences. In this work, we develop a multi-individual IBD inference method that is designed to handle discordant alleles in IBD segments, while still retaining the computational efficiency necessary to analyze biobank-scale data.

The new method retains important features of our previous method, such as the use of IBD transitivity to obtain linear scaling with sample size, and the application of a trim to the ends of pairwise IBD segments before applying transitivity to reduce false-positive IBD. Transitivity is the property that if haplotypes ℎ_1_and ℎ_2_ are IBD at a locus, and if haplotypes ℎ_2_ and ℎ_3_ are IBD at the locus, then haplotypes ℎ_1_ and ℎ_3_ must also be IBD. This is a natural property that multi-individual IBD should have, but the application of this property to inferred pairwise IBD segments can propagate false-positive errors in detected IBD. Thus, it is necessary to have a low rate of false positive error in the pairwise IBD segments that are used to infer multi-individual IBD. The endpoints of IBD segments tend to be difficult to determine accurately,^14^ so application of a trim results in a significant reduction in false positive IBD.

We provide a brief description of our new multi-individual IBD inference method here and provide further details in Section 1 of Supplemental Information.

We first apply a minor allele frequency (MAF) filter that excludes all markers whose second largest allele frequency is less than a threshold (0.1 by default). The MAF filter retains the most informative markers, reduces computation time, and reduces the number of discordant alleles in IBD segments.

We then identify a set of candidate pairwise IBD segments in the MAF-filtered data. The candidate-generating step uses four disjoint, interleaved sets of markers. The *k*-th marker (1 < *k* ≤ 4) set contains every fourth marker beginning with the *k*-th marker. The use of interleaved marker sets protects against loss of power due to discordant alleles, since a discordant allele will be present in only one of the four sets. We apply the Positional Burrows-Wheeler Transform (PBWT)^15^ to each marker set to identify identity-by-state (IBS) segments in the marker set that exceed a specified length (*L* = 1 cM, unless otherwise stated) and that are on adjacent haplotypes when the haplotype are lexicographically sorted by the sequence of alleles looking backwards from the last marker in the IBS segment.

For each pair of adjacent haplotypes with IBS segment length exceeding *L* cM, we use the ibd-ends algorithm with all markers that pass the MAF filter to estimate the endpoints of the underlying IBD segment.^14^ Each IBD segment endpoint is estimated as the median of the posterior endpoint distribution. The ibd-ends algorithm uses a probabilistic model that allows for mismatches that arise from genotype error, mutation, and gene conversion. The ibd-ends algorithm also accounts for inter-marker distances which can be large in regions with unmapped sequence reads, such as centromeres. Standard methods for detecting IBD segments based on IBS segment length can produce many false-positive IBD segments that span long inter-marker gaps.^16^ The ibd-ends algorithm accounts for these inter-marker gaps and does not have high false positive rates in these regions.^14^ We retain the IBD segment if the length estimated by the ibd-ends algorithm exceeds the *L* cM length threshold. We then trim *T* cM (*T* = 0.5 cM, unless otherwise stated) from each end of each IBD segment.

The preceding algorithm for identifying IBD segments will not find all IBD pairs since it only considers pairs of haplotypes that are adjacent when haplotypes are sorted by the PBWT. We fill in the missing IBD by enforcing IBD transitivity. When applying IBD transitivity, we take all the trimmed IBD segments that overlap a position and apply transitivity to define IBD haplotype clusters at that position.

### Gene conversion detection

We use the methodology described previously to detect alleles changed by gene conversion using multi-individual IBD.^10^ At each marker with MAF larger than a minimum, which is 0.1 in this work, we examine the IBD clusters at the position closest to the marker. We look for IBD clusters for which at least two haplotypes carry one allele and at least two haplotypes carry a different allele. These are the potential allele conversions (i.e., alleles changed by gene conversion). Each haplotype belongs to an individual, and that individual may be homozygous or heterozygous at the marker. If all the individuals with haplotypes in the cluster are homozygous at the marker, we do not record an allele conversion because the discordant alleles could be caused by the haplotypes in the IBD cluster carrying a cryptic deleted allele which results in the individuals carrying those haplotypes being called as homozygous for the individuals’ non-deleted alleles.

### Analysis of gene conversion

We estimate gene conversion rates in non-overlapping windows that have a fixed base pair length. In each window, we count the number of detected allele conversions. We divide the count by the expected heterozygosity, which is ∑_*i*_ 2 × *f*_*i*_ × (1 − *f*_*i*_) where *f*_*i*_ is the MAF of marker *i* and the sum is across the analyzed markers in the window (i.e., those passing the MAF filter). In homogeneous populations, the expected heterozygosity is proportional to the expected number of allele conversions, because alleles are only changed by gene conversion if the parent individual is heterozygous. In heterogenous populations, such as the TOPMed data analyzed in this study, the expected heterozygosity does not have this property but can still serve as a proxy for marker density. Our procedure provides relative rather than absolute rates of gene conversion, so we normalize the gene conversion rates. In the real human data, we normalize the rates to have mean 6 × 10^−6^ per bp across the autosomes,^4; 11^ while in the simulated data we normalize the rates so that the baseline simulations have mean 1 per bp in the region to facilitate comparison across different multiples of the baseline rate.

When calculating the correlation between an estimated gene conversion map and a crossover map, we ignore windows in which the IBD rate is more than 1.4 times or less than 0.6 times the median. The IBD rate at a locus is the proportion of pairs of haplotypes that are in the same IBD cluster, and the IBD rate for a window is the average IBD rate over loci in the window. A high IBD detection rate indicates natural selection.^14; 17; 18^ Natural selection leads to higher rates of IBD and hence a larger number of meioses in which gene conversions can be detected through IBD. Thus, natural section will tend to lead to increased gene conversion detection even if the gene conversion rate is not elevated. A low IBD detection rate can occur at chromosome ends, at centromeres and other regions devoid of genotypes, and in regions of high genotype error, and will lead to decreased gene conversion detection. Similarly, when calculating correlations between two estimated gene conversion maps, we ignore windows in which the IBD rate for either data set is more than 1.4 times or less than 0.6 times the median for that data set. We also ignore regions of low expected heterozygosity (i.e., low marker density) because these regions have less data and will have noisy results. If the expected heterozygosity is less than the window size divided by 10 kb (e.g. less than 1 for 10 kb windows or less than 100 for 1 Mb windows) we ignore the window. In the results with 10 kb, 100kb, and 1Mb windows, application of this heterozygosity filter removes 10%, 5%, and 2% respectively of the windows remaining after the application of the IBD rate filter.

### Simulated data

We simulate data from a growing population that is designed to be similar to modern human populations that have not gone through recent bottlenecks. The historical size of the population is 10,000 diploid individuals. The population has been growing at 3% per generation for the past 200 generations, for a current size of 3.7 million. Our simulation has a mutation rate of 1.5 × 10^−8^per bp per generation, a recombination rate of 10^−8^ per bp per generation, and gene conversions. Gene conversions have mean length 300 bp and a baseline initiation rate of 2 × 10^−8^ per bp per generation. Data simulated with msprime (see below) have gene conversion lengths following a geometric distribution, while data simulated with SLiM (see below) have gene conversion lengths distributed as a sum of two geometric random variables. As described below, some simulations include gene conversion hotspots with a higher rate of gene conversion, while other simulations have a higher rate of gene conversion across the entire simulated region. We add cryptic deletions to the data, in which individuals carrying the deletion allele are called as homozygous for their other allele. One percent of the simulated variants with frequency < 1% are turned into uncalled deletions with length drawn from an exponential distribution with mean 500 bp, and genotypes carrying the deletion are called as homozygous for the non-deleted allele.^10^ We add genotype error at rate 2 × 10^−4^. Genotypes affected by error have one of their alleles chosen at random to be changed. We phase the data using Beagle 5.4.^19^

We simulated 10,000 individuals across 20 regions of length 10 Mb using msprime v1.2 with the baseline level of gene conversion.^20; 21^ For these data, we generated ground-truth IBD and gene conversion information in order to calculate false discovery rates, using the methods described in our previous work.^10^

We simulated 125,000 individuals across 20 regions of length 10 Mb using msprime v1.2 with the baseline level of gene conversion (gene conversion tract initiation rate of 2 × 10^−8^ per bp), and a further 20 regions of length 10 Mb with 1.5 times the baseline level of gene conversion (gene conversion tract initiation rate of 3 × 10^−8^ per bp).

We also simulated data with gene conversion hotspots, for which we used SLiM v4 since msprime is limited to a constant gene conversion rate.^22; 23^ Our code for generating gene conversion hotspots follows suggestions in the SLiM manual and can be found in Section 2 of Supplementary Methods. We simulated the past 5000 generations with SLiM and then recapitated the simulations (added further generations as needed to complete coalescence) and added mutations using pyslim and msprime.^24^ We simulated 125,000 individuals across 10 regions of length 10 Mb, with the central 10 kb of those regions having twice the baseline gene conversion rate, and a further 10 regions with the same parameters except for a ten-fold rather than two-fold hotspot gene conversion rate.

### PRDM9 binding enrichment

We used published PRDM9 binding enrichment scores that were obtained from expressing PRDM9 in a human cell line and performing ChIP-seq to assess binding (see Web resources).^25^ The data that consisted of 170,198 PRDM9 binding peaks across the genome. We lifted the positions over from hg19 to GRCh38 to match the sequence data described below. When partitioning and analyzing the data in windows, we summed the enrichment scores for peaks having their centers in each window.

### TOPMed data

We analyzed phased whole autosome sequence data from a previous phasing of 39,961 TOPMed individuals,^19^ but with 1882 individuals from the withdrawn SARP study removed, with a resulting size of 38,079 individuals. The analyzed individuals are multi-ethnic with a predominance of European ancestry and are mostly from the USA.^26^

### UK Biobank data

We analyzed phased whole autosome data on 125,361 individuals of White British ancestry from a previous phasing of UK Biobank individuals.^27; 28^

## Results

### Length and trim parameter settings

Using the simulated data on 10,000 individuals that have true IBD and gene conversion information, we calculated detection rates and false discovery rates for IBD and allele conversions (Table S1). IBD false discovery rates are less than 1% and allele conversion false discovery rates are less than 2% for all settings with *T* ≥ 0.75. IBD and allele conversion false discovery rates are both less than 3% for all settings with *T* ≥ 0.5.

Using the simulated data with 125,000 individuals with a constant gene conversion rate of 1.5 times the baseline rate, we investigated the level of bias in estimation of the relative gene conversion rate (Table S2). We find that there is a small downward bias when the gene conversion rate is high, but that for *T* ≥ 0.5 the bias is small, with the estimated relative rate being 1.48 while the true relative rate is 1.5. Increasing the length threshold (*L*) to values larger than 1 has little effect on bias, and it reduces the number of detected allele conversions (Table S1).

We then investigated which length and trim settings give the highest accuracy when estimating the gene conversion rate in the TOPMed and UK Biobank data. The primary metric that we use is Pearson’s correlation between the estimated gene conversion rate and the sex-averaged crossover rate. Previous work has shown that the gene conversion rate tends to be high in regions where the recombination rate is high,^2; 6^ so a higher correlation indicates more accurate estimation of gene conversion rates. As a secondary metric, we consider the correlation between the gene conversion rate estimates from the two data sets. Although there may be some population differences in the maps, we expect them to be similar.

We found that *L* = 1 with *T* = 0.5 gave the best or close to best results on these metrics across the two data sets when considering windows of size 10 kb and of size 1 Mb (Tables S3 and S4). Since these parameter settings were also supported by the simulated data, we use these settings for subsequent analyses.

### Detection of hotspots and inter-window differences in gene conversion rates

We used simulated data with gene conversion hotspots to investigate the power to detect gene conversion hotspots, and we used simulated data with baseline and 1.5x baseline gene conversion rates to investigate the accuracy of estimated gene conversion rates in 100 kb and 1 Mb windows.

Using the simulated data with 125,000 individuals and gene conversion hotspots, we investigated the ability to estimate the relative rate of gene conversion in 10kb windows. We removed windows within 1 cM of each end of the analyzed region before presenting results, because IBD rates are zero or significantly reduced in these end regions.

Figure 1A shows that the median estimates for hotspots with twice (2x) and ten times (10x) the baseline gene conversion rate are close to their true values. There is some overlap in the distribution of estimates from baseline gene conversion rate windows compared with estimates from windows with twice the baseline gene conversion rate, however 100% (10/10) of the 2x estimates exceed the 99^th^ percentile of the baseline distribution. Estimated gene conversion rates in hotspots with 10x gene conversion rate are completely separated from both the baseline and the 2x gene conversion rates. An increase in downward bias is seen in Figure 1A as the hotspot intensity increases. High gene conversion rates can reduce IBD detection power, both directly due to creating allele mismatches in IBD segments and indirectly through the impact of these mismatches on haplotype phasing accuracy.

**Figure 1:**
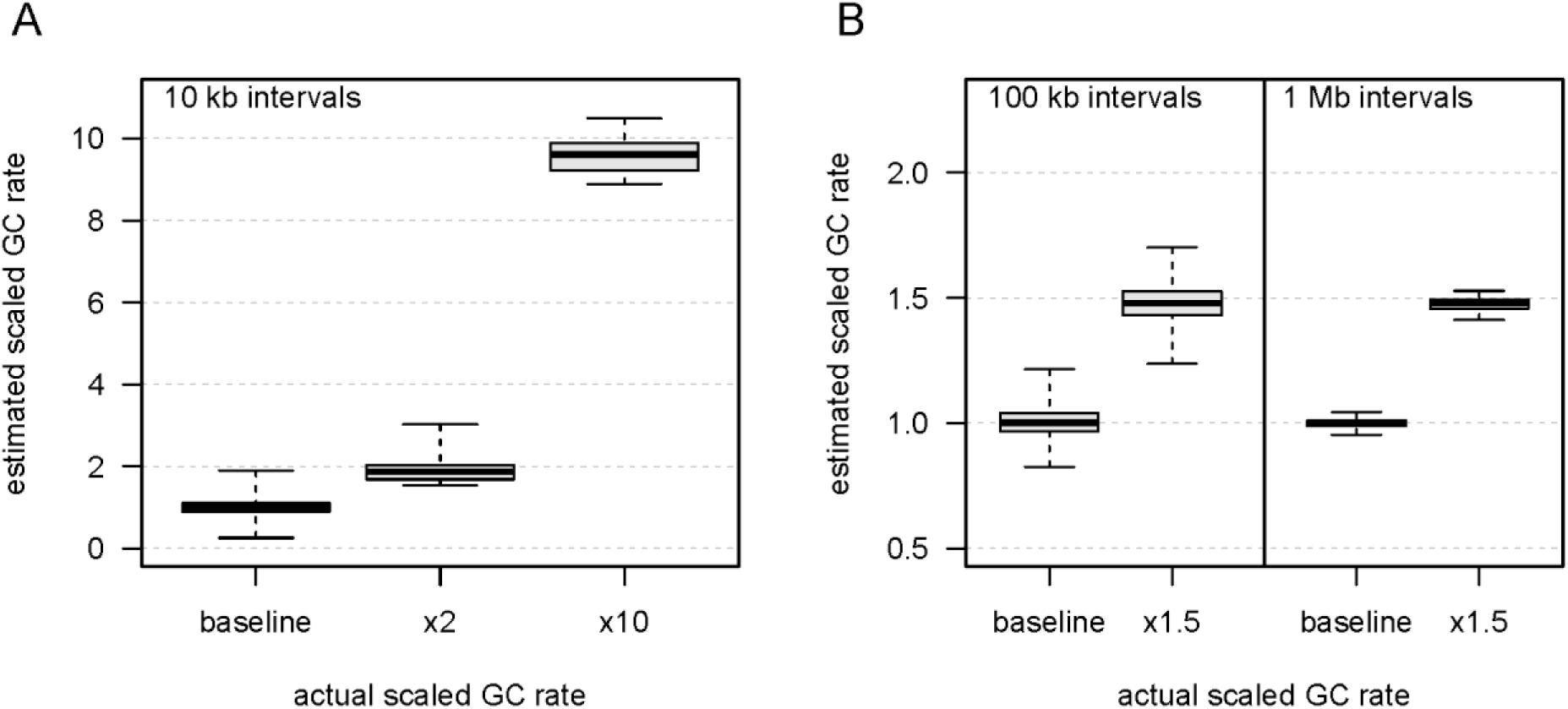
Estimated scaled gene conversion rate in simulated data. Simulated data have 125,000 individuals. Boxplots show range, interquartile range, and median. All analyses use a *L* = 1 cM IBD length threshold and a *T* = 0.5 cM end trim. A. 10 kb hotspots with twice (x2) or ten times (x10) the baseline gene conversion rate. We estimate relative gene conversion rates in 10 kb windows and scale them so that the mean baseline rate is 1. Baseline results are based on 15,678 windows with heterozygosity sum greater than 1 and located at least 1 cM from the ends of the simulated regions, while hotspot results are each based on 10 windows (one window from each of 10 simulated regions). B. 100 kb and 1 Mb regions with baseline gene conversion or 1.5 times (x1.5) the baseline rate. We estimate relative gene conversion rates in 100kb or 1 Mb windows and scale them so that the mean baseline rate for the 1 Mb windows is 1. The 100 kb boxplots are each based on 1600 windows located at least 1 cM from the ends of the simulated regions, while the 1 Mb boxplots are each based on 160 windows located at least 1 cM from the ends of the simulated regions.

Using the simulated data with 125,000 individuals and constant gene conversion rate, we investigated the ability to estimate gene conversion rates in long windows. Figure 1B shows that with 100 kb or 1 Mb windows, there is no overlap between the baseline and 1.5x results, so there is very high power to distinguish a 1.5 factor difference from baseline.

### Gene conversion maps from TOPMed and UK Biobank autosome data

We detected 3,503,072 allele conversions in the TOPMed data and 13,901,830 allele conversions in the UK Biobank data. For comparison, an analysis of the UK Biobank data with our previous multi-individual IBD detection method found 9,313,066 allele conversions. We collated the allele conversions into non-overlapping windows of length 10 kb, 100 kb, or 1 Mb, and we estimated the gene conversion rate in each window as described in Methods. We removed from further analysis any window that had an IBD rate more than 40% higher or 40% lower than the median IBD rate in either data set. We also removed windows with expected heterozygosity less than the required minimum in either data set (100 for the 1 Mb windows, 10 for the 100 kb windows, and 1 for the 10 kb windows).

In humans, gene conversion hotspots tend to co-occur with crossover hotspots.^2; 4^ Table 1 shows the correlation between our two gene conversion rate maps (TOPMed and UK Biobank) and the deCODE sex-averaged crossover map,^29^ as well as the correlation between our two gene conversion maps. Correlations increase with increasing window size because larger windows contain more data and thus have higher relative accuracy. At a 1 Mb resolution, our TOPMed gene conversion rate map has a correlation of 0.667 with the deCODE sex-averaged crossover map. For comparison, we averaged the maternal and paternal non-crossover (gene conversion) deCODE maps,^6^ which are based on overlapping 3 Mb windows, and found a correlation of 0.553 with the deCODE sex-averaged crossover map. This reduced correlation for the deCODE gene-conversion map compared to our TOPMed gene conversion map may be due to our TOPMed map being based on more than 55 times more observed allele conversions than the deCODE map.

**Table 1:** Pearson correlation coefficients between the sex-averaged deCODE 2019 crossover map and our inferred gene conversion maps based on TOPMed and UK Biobank data for 10 kb, 100 kb, and 1 Mb windows.

| Window | TOPMed vs<br>deCODE | UK Biobank vs<br>deCODE | TOPMed vs<br>UK Biobank |
| --- | --- | --- | --- |
| 10 kb | 0.561 | 0.426 | 0.632 |
| 100 kb | 0.597 | 0.457 | 0.665 |
| 1 Mb | 0.667 | 0.626 | 0.855 |

Our UK Biobank gene conversion rate map has lower correlation than our TOPMed gene conversion rate map with the deCODE crossover map, especially at finer scales of resolution (10 kb or 100 kb). When we restrict the analysis to windows in which our two gene conversion rate maps are similar, we find that correlations with the crossover map increase slightly for our TOPMed map and increase significantly for our UK Biobank map, so that the UK Biobank correlations become similar to the TOPMed correlations (Table S5). This suggests that our UK Biobank map contains more artifacts than our TOPMed map.

Figure 2 shows estimated gene conversion rates from the TOPMed data along the autosome for 1 Mb windows. The estimated rates of gene conversion are elevated near the chromosome ends. In males, crossover recombination occurs at greatly elevated rates in the subtelomeric regions,^30^ thus leading to high sex-averaged crossover rates in these regions (Figure S1), so it is not surprising to see this effect for gene conversion recombination as well. Our TOPMed gene conversion maps for 10 kb, 100 kb, and 1 Mb windows are provided as Supplemental Information.

**Figure 2:**
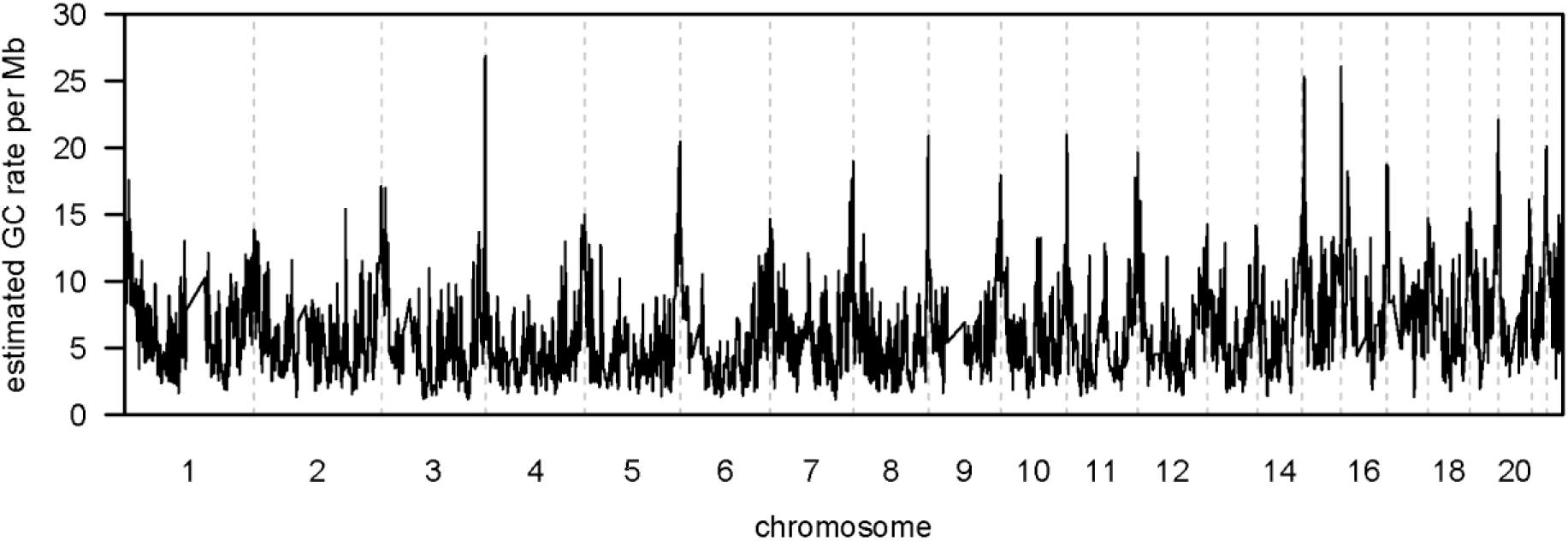
Estimated gene conversion rates for TOPMed data in 1 Mb windows across the autosomes. Estimated relative gene conversion rates have been scaled to have mean 6 per Mb (6 × 10^−6^ per base pair). Estimates are calculated in 1 Mb windows, and windows with expected heterozygosity less than 100 or IBD rate more than 40% higher or 40% lower than the median IBD rate are excluded. Chromosomes are separated by vertical gray dashed lines.

We plotted gene conversion rates in the vicinity of the strongest gene conversion hotspots (Figure 3 and Figure S2). These figures show that hotspot peaks die away over very short distances, typically within 1 kb.

**Figure 3:**
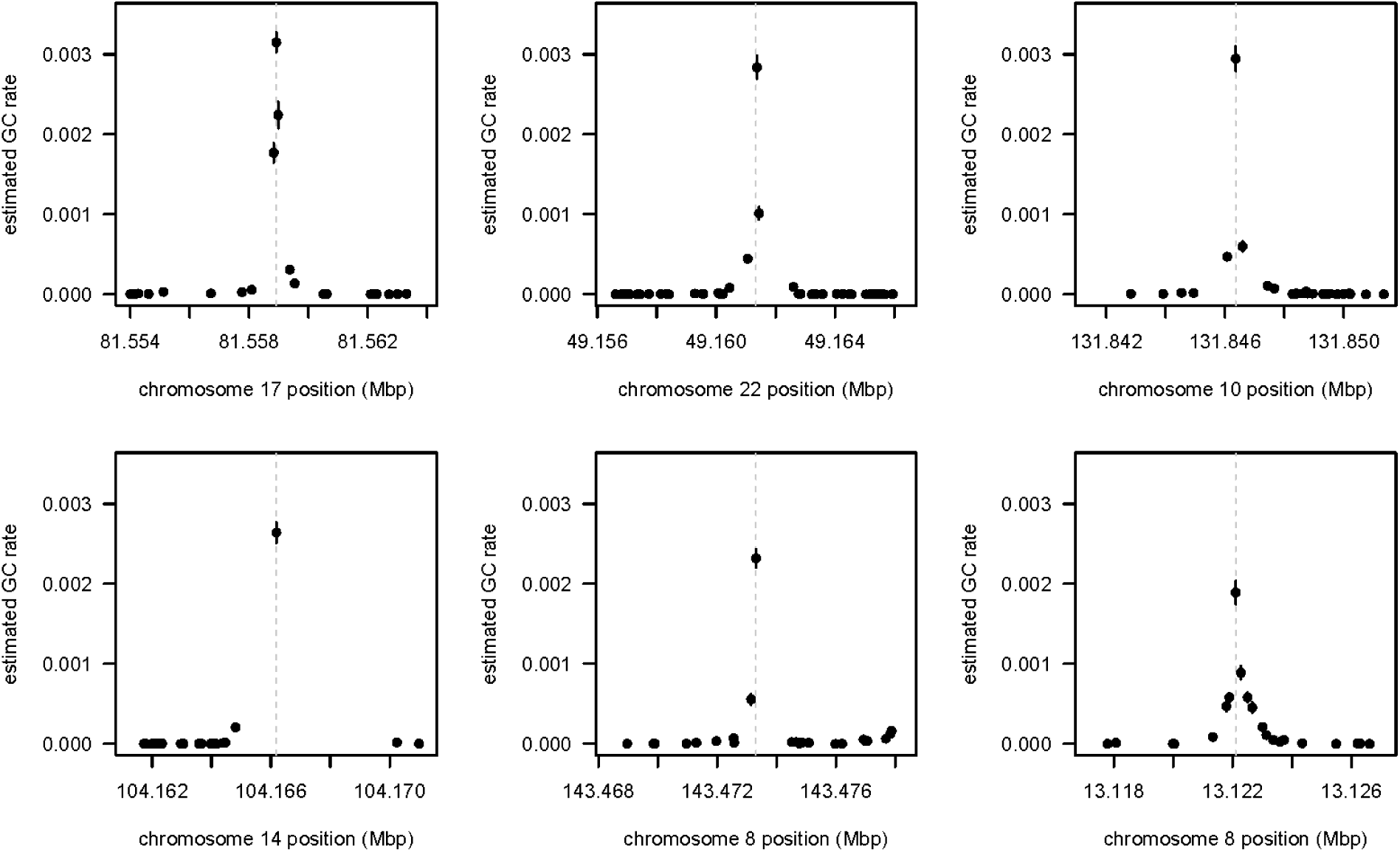
Gene conversion hotspots. We selected the markers with the highest estimated gene conversion rates in the TOPMed data and plotted the gene conversion rates in the UK Biobank data at nearby markers. The gray dashed vertical lines give the locations of the hotspots in the TOPMed data. TOPMed hotspots that are within 10 kb of a marker with a larger gene conversion rate are omitted, as are hotspots at markers for which the TOPMed IBD rate is more than 1.4 times or less than 0.6 times the median. TOPMed hotspots for which there is no UK Biobank marker with MAF > 10% within 100 bp or for which there are fewer than 20 UK Biobank markers with MAF > 10% within the 10 kb region centered on the TOPMed hotspot position are also omitted because the plots do not show sufficient detail. Plots are shown in order of TOPMed hotspot rate with highest first, left to right, then top to bottom. The estimated gene conversion (GC) rate at a marker (in the TOPMed data to select the hotspots, and in the UK Biobank data for the y-axis values in these plots) is the number of detected allele conversions divided by the expected heterozygosity of the marker, normalized so that the autosome-wide average is 6 × 10^−6^. Estimates are plotted as dots, while 95% confidence intervals are given as vertical lines through the dots and are obtained by assuming that the number of detected allele conversions follows a Poisson distribution to obtain the standard error and then adding two standard errors to each side of the estimate. Each plot shows all UK Biobank markers with MAF > 10% within 5 kb on either side of the hotspot location. Positions on the x-axes are in GRCh38 coordinates. This figure shows the top six hotspots meeting the UK Biobank marker density criteria, and Figure S2 shows the top twenty-four such hotspots.

### PRDM9 binding enrichment

One question of interest is whether PRDM9 binding enrichment is more strongly predictive of gene conversion or of crossovers. To answer this question, we estimated the correlation of PRDM9 binding enrichment (see Methods) with our gene conversion rate estimates from the TOPMed data and with the deCODE sex-averaged crossover rates. For windows of size 100 kb or 1 Mb, we find higher correlation between PRDM9 binding enrichment and the gene conversion rate than between PRDM9 binding enrichment and the crossover rate (Figure 4). For example, with 1 Mb windows, the Pearson correlation coefficient is 0.52 for gene conversion and 0.34 for crossovers.

**Figure 4:**
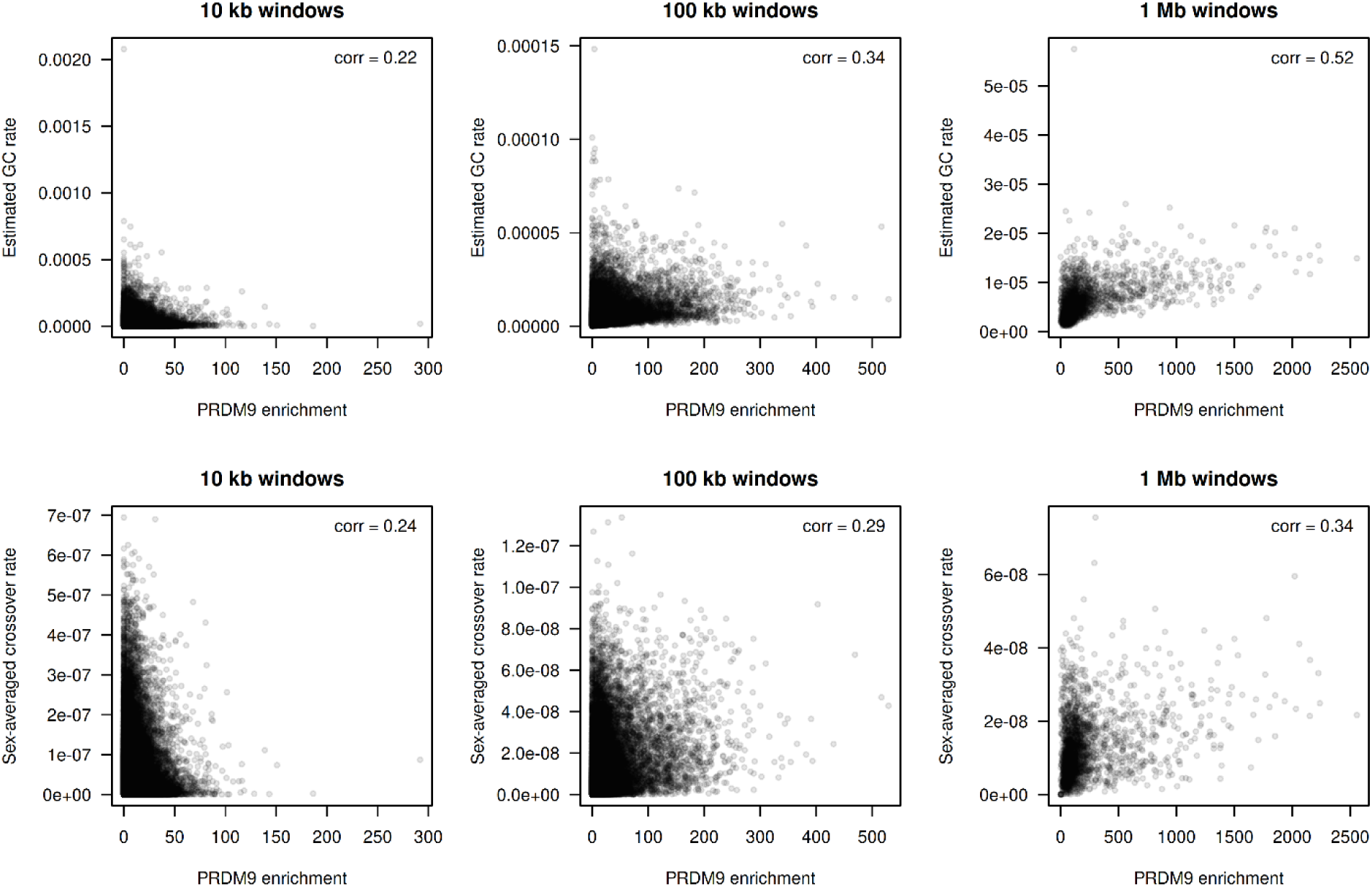
Comparison of gene conversion rate estimates from the TOPMed data and sex-averaged crossover rates with PRDM9 binding enrichment. Gene conversion rates are shown in the upper row, while crossover rates are shown in the lower row. Each column has a different window size which is notated above the plots. The Pearson correlation coefficient between the gene conversion rate or crossover rate and PRDM9 binding enrichment is shown in the upper right of each plot.

For 10 kb windows, crossover rates have a higher correlation than gene conversion rates with PRDM9 binding enrichment (0.24 for crossovers and 0.22 for gene conversion). The total number of crossovers used to build the crossover map is 4.5 million, or approximately 15 crossovers per 10 kb on average.^29^ The total number of allele conversions used to build our TOPMed gene conversion map is 3.5 million, or approximately 11 allele conversions per 10 kb on average. The lower number of allele conversion events compared to crossover events combined with the fact that the variance will be high relative to the mean at the 10 kb scale due to the low average numbers of events in 10 kb windows may be the primary factor underlying the lower correlation of the gene conversion map at this scale.

We used enrichment peak centers as the locations for this analysis; the PRDM9 binding map also provides confidence intervals for the peak locations. These have a median length of 55 bp (in hg19 coordinates), which is much shorter than the window sizes that we are considering.

### Computing times

Inferring multi-individual IBD on chromosome 1 with *L* = 1 and *T* = 0.5 took 73 minutes on a 24-core compute node for the 38,079-individual TOPMed data and 191 minutes on a 96-core compute node for the 125,361-individual UK Biobank data.

## Discussion

We presented a new method for multi-individual IBD detection and applied it to detecting allele conversions and estimating gene conversion rates in 10 kb, 100 kb, and 1 Mb windows.

The first stage of our multi-individual IBD detection method generates a candidate set of pairwise IBD segments that are evaluated in the second stage using the ibd-ends probabilistic model. The challenge of detecting IBD segments in the presence of discordant alleles caused by mutation, gene conversion, and genotype error is addressed in the first stage by performing IBS segment detection separately on four disjoint, interleaved marker sets and in the second stage with a probabilistic model that allows for discordant alleles. The probabilistic ibd-ends algorithm also allows our algorithm to model uncertainty in IBD segment endpoints. Our two-stage method avoids the problem of quadratic scaling of pairwise IBD segment detection with sample size through the use of the Positional Burrows-Wheeler Transform and IBD transitivity. The result is an algorithm that scales linearly with sample size in both computing time and output file size.

We generated gene conversion rate maps using both UK Biobank data, and TOPMed data. Although the UK Biobank data contained 3.3 times as many individuals and resulted in detection of almost 4 times as many allele conversions, we found that the map generated from the TOPMed data had higher correlation with the deCODE crossover map, which suggests that the TOPMed map is superior. This difference may be due to differences in the sequencing and QC pipelines between the two data sets. We also found that our TOPMed gene conversion map was more highly correlated than the deCODE gene conversion map with the deCODE crossover map, which suggests that the TOPMed gene conversion map is more accurate than the deCODE gene conversion map. Our TOPMed map is sex-averaged and has reasonable accuracy in 10 kb windows, whereas the deCODE gene conversion maps are sex-specific and use 3 Mb windows. Our TOPMed-based gene conversion map has an average of 1.1 allele conversions per kb. In contrast, the deCODE gene conversion maps have an average of around 20 allele conversions per Mb combined across both sexes.

At scales of 100 kb and 1 Mb, our TOPMed-based gene conversion rate map was more highly correlated than the deCODE crossover map with a map of PRDM9 binding enrichment. Since the deCODE crossover map has high resolution at these scales and is expected to be highly accurate due to its pedigree-based design, this suggests that PRDM9 binding has a stronger local effect on gene conversion than on crossing-over.

The most direct way to estimate gene conversion rates is to observe products of meioses, such as through sperm-typing or family data. In order to achieve highest accuracy with this type of approach, it is necessary to have multi-generational families.^4^ Even with very large data, such as the recently published deCODE gene conversion data with 10,840 meioses, the number of observed events (62,762 in the deCODE gene conversion data) is not sufficient for obtaining a high-resolution map. An indirect way to estimate gene conversion rates is to construct LD-based gene conversion maps in a similar way to the construction of LD-based crossover maps.^7; 8^ However, these LD-based maps also have low resolution.^9^ Our IBD-based method has both similarities and differences with these alternate approaches. Unlike the LD-based approaches, but like the family-based approaches, we observe specific allele conversions, and like the multi-generational family-based approaches, we have excellent control over false positive observations. However, compared to family-based approaches we observed orders of magnitude more events, allowing for good resolution even at a 10 kb scale with our TOPMed map.

A disadvantage of our IBD-based approach compared to family-based approaches is that we cannot assign observed allele conversions to specific meioses. Thus, whereas the recently published deCODE study was able to estimate sex-specific gene conversion rates, age effects, and genetic associations between genome-wide gene conversion rates and specific alleles, we cannot estimate these quantities with our method. On the other hand, because we observe a large number of events, we are able to visualize the decay of events around hotspots and to observe a higher correlation of gene conversion than crossovers with PRDM9 binding.

## Supporting information

Gene conversion map with 1 Mb windows

Gene conversion map with 10 kb windows

Gene conversion map with 100 kb windows

## Acknowledgements

The methodological and analytical work performed in this study was supported by the National Human Genome Research Institute (NHGRI) under award numbers R01 HG005701 and R01 HG008359. This research has been conducted using the UK Biobank Resource under Application Number 19934. The content is solely the responsibility of the authors and does not necessarily represent the official views of the National Institutes of Health or the UK Biobank.

Sequence data for the Trans-Omics in Precision Medicine (TOPMed) program was supported by the National Heart, Lung and Blood Institute (NHLBI). Core support including centralized genomic read mapping and genotype calling, along with variant quality metrics and filtering were provided by the TOPMed Informatics Research Center (3R01HL-117626-02S1; contract HHSN268201800002I). Core support including phenotype harmonization, data management, sample-identity QC, and general program coordination were provided by the TOPMed Data Coordinating Center (R01HL-120393; U01HL-120393; contract HHSN268201800001I). We gratefully acknowledge the studies and participants who provided biological samples and data for TOPMed. Funding for the Barbados Asthma Genetics Study was provided by National Institutes of Health (NIH) R01HL104608, R01HL087699, and HL104608 S1. The Framingham Heart Study was supported by contracts NO1-HC-25195, HHSN268201500001I and 75N92019D00031 from the NHLBI and grant supplement R01 HL092577-06S1; genome sequencing was funded by HHSN268201600034I and U54HG003067. See Supplemental Data for acknowledgments of additional studies in the TOPMed data.

## Author contributions

BLB developed the IBD haplotype clustering method and software; SRB. developed the method for estimating gene conversion rates; SRB designed and performed the analyses; SRB and BLB wrote the manuscript.

## Declaration of interests

The authors declare no competing interests.

## Web resources

### ibd-cluster program (version 0.2)

https://github.com/browning-lab/ibd-cluster

### PRDM9 binding peaks

https://www.ncbi.nlm.nih.gov/geo/download/?acc=GSE99407&format=file&file=GSE99407_ChIPseq_Peaks.YFP_HumanPRDM9.antiGFP.protocolN.p10e-5.sep250.Annotated.txt.gz

(accessed August 8, 2024).

### TopMed sequence data

https://topmed.nhlbi.nih.gov/topmed-data-access-scientific-community

### UK Biobank sequence data

https://www.ukbiobank.ac.uk/

## Data and code availability

The TopMed and UK Biobank data sets analyzed in this study are available to researchers upon approval of a data access application. The open source ibd-cluster software and gene conversion rate estimates are publicly available (see Web resources and Supplemental Information).

## Notes

### Competing Interest Statement

The authors have declared no competing interest.

