## Supplemental Information for "Estimating gene conversion rates from population data using multi-individual identity by descent"

#### 1 Inference of multi-individual identity by descent

##### 1.1 Method overview

**Definitions**: A pairwise haplotype segment is a pair of haplotypes that are restricted to a chromosome segment. A pairwise haplotype segment is an identity-by-state (IBS) segment if the two haplotypes have identical alleles in the segment. A pairwise haplotype segment is an identity-by-descent (IBD) segment if both haplotypes have inherited the segment identically by descent from a common ancestor.

We use the Positional Burrows-Wheeler Transform (PBWT)[1] to identify a set of pairwise haplotype segments that have a long subsequence of shared alleles. We then apply the ibd-ends algorithm[2] to estimate the endpoints of the IBD segment containing the midpoint of each pairwise haplotype segment. If the length of the estimated IBD segment is $\geq L$ cM, we trim $T$ cM from each end of the IBD segment and store the trimmed segment. The default minimum IBD segment length is $L = 1$ cM, and the default end trim is $T = 0.5$ cM.

We use the set of trimmed IBD segments to cluster IBD haplotypes. Two haplotypes $h_1$ and $h_n$ are assigned to the same IBD cluster at a locus if there is a sequence of haplotypes $h_1, h_2, \ldots, h_n$ such that each pair of consecutive haplotypes $(h_i, h_{i+1})$ has a trimmed IBD segment that contains the locus.

Our IBD haplotype clustering algorithm scales linearly with sample size because the PBWT, the number of pairwise haplotype segments that we analyze, the IBD haplotype clustering, and the output file size scale linearly with sample size. The following sections describe each step of our method.

#### 1.2 Identifying a set of candidate pairwise IBD segments

We first exclude any marker whose second largest allele frequency is less than a user-specified threshold (0.1 by default). The allele frequency filter reduces computation time, memory requirements, and the number of discordant alleles in IBD segments.

We then index the remaining markers in chromosome order with $1, 2, \ldots, M$, and we partition the $M$ markers into four interleaved sets. The $k$-th set contains every 4th marker beginning with marker $k$. We retain the original marker indices, so the indices of the markers in the $k$-th set are $k, k+4, k+8$, etc. The use of four interleaved marker sets increases power to detect long IBD segments that contain discordant alleles caused by genotype error, mutation, or gene conversion.

We then apply the Positional Burrows-Wheeler Transform (PBWT)[1] to each marker set. At each marker $m$, the PBWT defines a positional prefix array $a_m$ and a divergence array $d_m$.[1] The positional prefix array $a_m$ is the sorted list of haplotype indices when haplotypes are lexicographically sorted by their reverse prefix. A haplotype's reverse prefix at marker $m$ is the sequence of preceding alleles in reverse order. The $(j+1)$-st element of the divergence array $d_m$ is the first marker in the IBS segment containing marker $m-1$ for haplotypes $a_m[j]$ and $a_m[j+1]$ in the positional prefix array. Durbin's Algorithm 2 gives pseudocode for computing $a_{m+1}$ and $d_{m+1}$ from $a_m$ and $d_m$.[1] Since we apply the PBWT to marker sets that contain every 4-th marker, we modify Durbin's algorithm 2 to compute $a_{m+4}$ and $d_{m+4}$ from $a_m$ and $d_m$.

If marker $m$ is in the $k$-th marker set, the haplotype that shares the longest preceding allele sequence in the $k$-th marker set with haplotype $a_m[j]$ will be either $a_m[j-1]$ or $a_m[j+1]$ due to the reverse prefix sorting. For each marker $m$, we check each pair of consecutive haplotypes $a_m[j]$ and $a_m[j+1]$ in the positional prefix array to determine if the pair have an IBS segment that ends with the preceding marker in the set and whose length is $\geq S$ cM ($S = 1$ cM by default). This is equivalent to checking whether haplotypes $a_m[j]$ and $a_m[j+1]$ carry different alleles at marker $m$ and whether the distance between marker $d_m[j+1]$ and marker $(m-4)$ is $\geq S$ cM. If the IBS segment length is $\geq S$, the pairwise IBS haplotype segment is stored. The stored pairwise haplotype segment is an IBS segment in the $k$-th market subset, but it is not necessarily an IBS segment in the set of all $M$ markers.

The maximum number of stored pairwise haplotype segments scales linearly with sample size because each pair of haplotypes is adjacent in the positional prefix array $a_m$ for some marker $m$ and because the length of each segment is $\geq S$ cM. Thus, if a chromosome has length $C$ cM and if there are $H$ haplotypes and 4 interleaved marker sets, the maximum number of stored pairwise haplotype segments is $4H(C/S)$. This linear scaling makes it possible to apply our method to data sets with millions of individuals.

From here on, we consider the stored pairwise haplotype segments as segments in the set of all $M$ markers. We merge overlapping pairwise haplotype segments from the four PBWT analyses. If two pairwise haplotype segments have the same pair of haplotypes and have overlapping chromosome intervals, the two pairwise haplotype segments are replaced by a merged segment whose chromosome interval is the union of the overlapping intervals.

#### 1.3 Estimating IBD segment endpoints

After merging overlapping pairwise haplotype segments, we apply the ibd-ends algorithm to estimate the endpoints of the IBD segment that contains the midpoint of each pairwise haplotype segment.[2] The ibd-ends probabilistic model is constructed from the phased genotypes for all individuals and a genetic map, and it allows for discordant alleles in IBD segments. The ibd-ends method estimates a probability distribution for each IBD segment endpoint, and we set the endpoint to be the 50-th percentile of the endpoint distribution.

#### 1.4 IBD clustering

After we estimate the IBD segments' endpoints, we exclude IBD segments with length $< L$ cM, and we trim $T$ cM from each end of each remaining IBD segment. We use the trimmed segments to cluster haplotypes at genetic map positions that are multiples of $D$ cM ($D, 2D, 3D, \ldots$). $D = 0.02$ cM by default.

We assign two haplotypes to the same IBD cluster at a position if the two haplotypes have a trimmed IBD segment that overlaps the position. Haplotype clustering at a marker is carried out efficiently via a disjoint set data structure that represents the haplotypes in an IBD cluster as a tree. We begin with one haplotype per cluster and each cluster represented as a singleton tree. If a trimmed IBD segment contains the locus, we merge the trees (i.e. IBD clusters) containing the two haplotypes in the IBD segment. We perform path compression and union by rank when merging trees to increase computational efficiency.[3]

#### 1.5 Output

After performing IBD clustering at a locus, we index the IBD haplotype clusters at the locus. If there are $H$ haplotypes, and $I$ clustering loci, the output contains an $I \times H$ matrix of cluster indices. The $(j, k)$-th element of the output matrix is the cluster index for the $k$-th haplotype in the input VCF file at the $j$-th clustering locus.

### 2 Simulation with variable gene conversion rate

The following SLiM v4 code is for simulating gene conversion hotspots. The implementation of the hotspots can be found in the "recombination" section of the code. This code outputs tree sequences, that can then be input to tskit/msprime for recapitation, addition of mutations, and output of a vcf file.

```
// Output sample size is NSAMP diploid individuals
// Region size is REGSIZE
// Gene conversion is included, and has a hotspot
// input variables are HSIZE, HRATIO, NSAMP, REGSIZE, SETTING
// HRATIO should be > 1, and HSIZE should be less than REGSIZE
// SETTING is for labeling the output
// These commands are written in slim 4
// Example for use follows on next 3 lines ("slim_growth" is the name of this
file of slim commands):
// hsize=10000; hratio=2; nsamp=125000; regsize=10000000; setting=3
// SLiM4/slim -d HSIZE=$hsize -d HRATIO=$hratio -d NSAMP=$nsamp \
// -d REGSIZE=$regsize -d SETTING=${setting} -s $seed slim_growth

initialize() {
      initializeTreeSeq(); // turn on the tree sequence recording
      initializeMutationRate(0);
      initializeMutationType("m1", 0.5, "f", 0.0);
      initializeGenomicElementType("g1", m1, 1.0);
      chromend = REGSIZE - 1;
      initializeGenomicElement(g1, 0, chromend);
      gcprop = 2.0/3.0; // this proportion of crossovers will result in gc
      gcmeanlen = 300; // mean gc length is 100 bp
      modrecomb = 1e-8/(1-gcprop); // more crossovers since some will be gcs.
      modrecomb2 = HRATIO*modrecomb;
      prestarthot = asInteger(REGSIZE/2-HSIZE/2-1);
      endhot = asInteger(REGSIZE/2+HSIZE/2-1);
      initializeRecombinationRate(c(modrecomb,modrecomb2,modrecomb),
            c(prestathot,endhot,chromend));
      // added a hotspot (affects both crossover and gc, but we'll take out
      // the extra crossovers), starts after prestarthot and ends at endhot
      initializeGeneConversion(gcprop,gcmeanlen,1.0);
}
```

```
recombination() {
      // remove some proportion of single crossovers in the hotspot region
      // need to keep all double crossovers (gcs), i.e. those within 3 kb
      prestarthot = asInteger(REGSIZE/2-HSIZE/2-1);
      endhot = asInteger(REGSIZE/2+HSIZE/2-1);
      retainprop = 1.0/HRATIO;
      nbreaks = size(breakpoints);
      if(nbreaks==0) return F;
      if(nbreaks>1){
        isdoublebreak = c();
        for(i in 1:nbreaks){
          thismin = 5000;
          for(j in 1:nbreaks){
           if(i==j) next;
           dist = abs(breakpoints[i-1]-breakpoints[j-1]);
           if(dist < thismin) thismin = dist;
           }
         if(thismin < 3000) isdoublebreak = c(isdoublebreak,T);
         else isdoublebreak = c(isdoublebreak,F);
         }
        }
       else isdoublebreak = c(F);
       outregion = breakpoints <= prestarthot | breakpoints > endhot;
       retain = outregion | runif(nbreaks) < retainprop;
       retain2 = retain | isdoublebreak;
       if(!all(retain2)){
         breakpoints = breakpoints[retain2];
         return T;
         }
       else
        return F;
}
////// apply the demographic model model
// Start with out of Africa bottleneck
1 early() { sim.addSubpop("p1",10000); }
// Growth starting 200 generations ago, 3%
99801:100000 early() {
newSize = 10000*exp(log(1.03)*(sim.cycle-99800));
 p1.setSubpopulationSize(asInteger(newSize));
}
// Reduce size to simulate sampling of individuals from the population
100001 early() {
 newSize = NSAMP;
 p1.setSubpopulationSize(newSize);
 }
// output data
100001 late()
{
 sim.treeSeqOutput("slim_data/setting"+SETTING+"_seed"+getSeed()+".trees");
}
```

### 3 Supplemental Figures

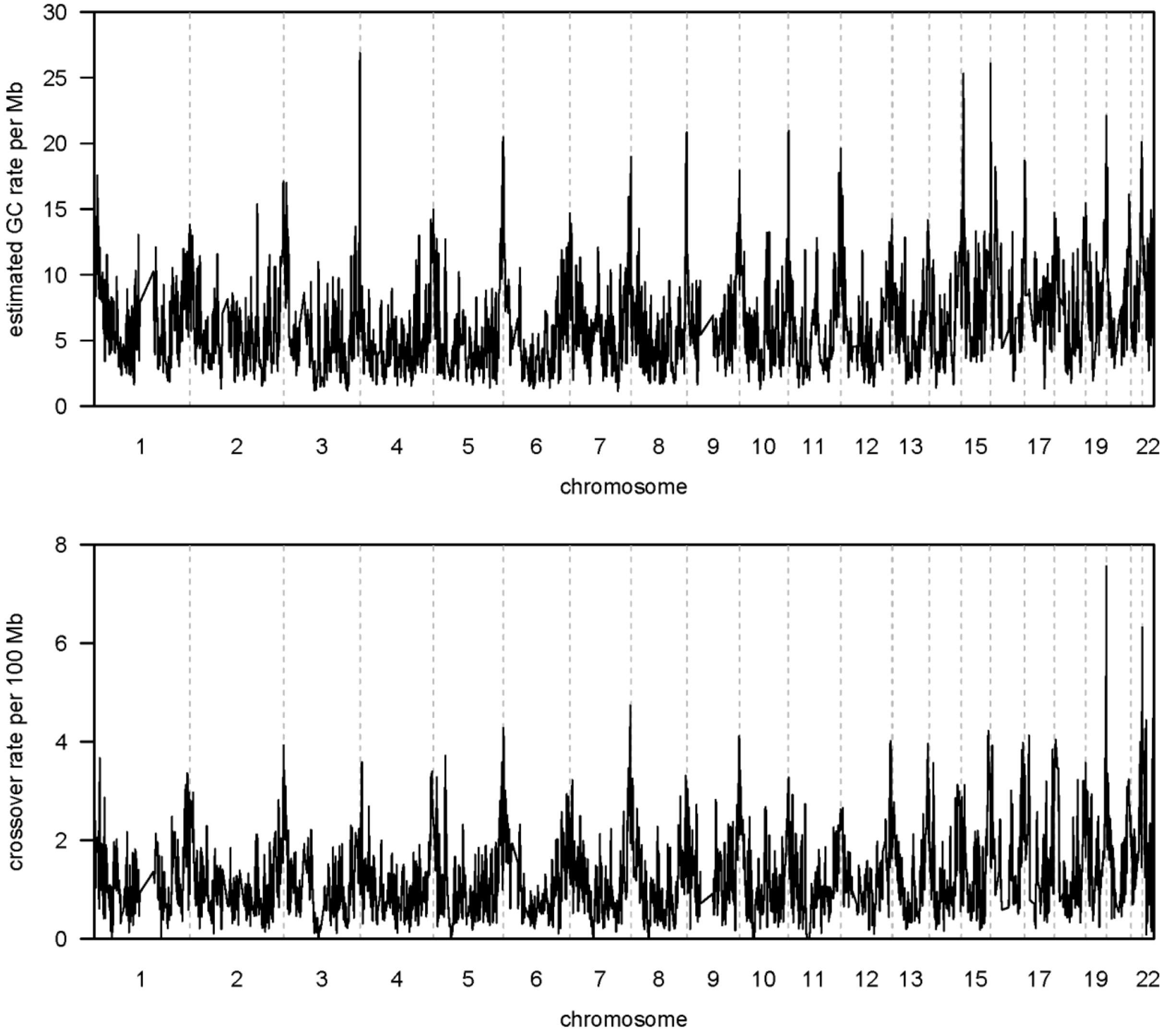


**Figure S1: Estimated gene conversion rates and crossover rates in 1 Mb windows across the autosomes.** The estimated gene conversion rate from the TOPMed data is shown in the top panel. Estimated relative gene conversion rates have been scaled to have mean 6 per Mb ($6 \times 10^{-6}$ per base pair). The sex-averaged crossover rate from the deCODE 2019 map is shown in the bottom panel.

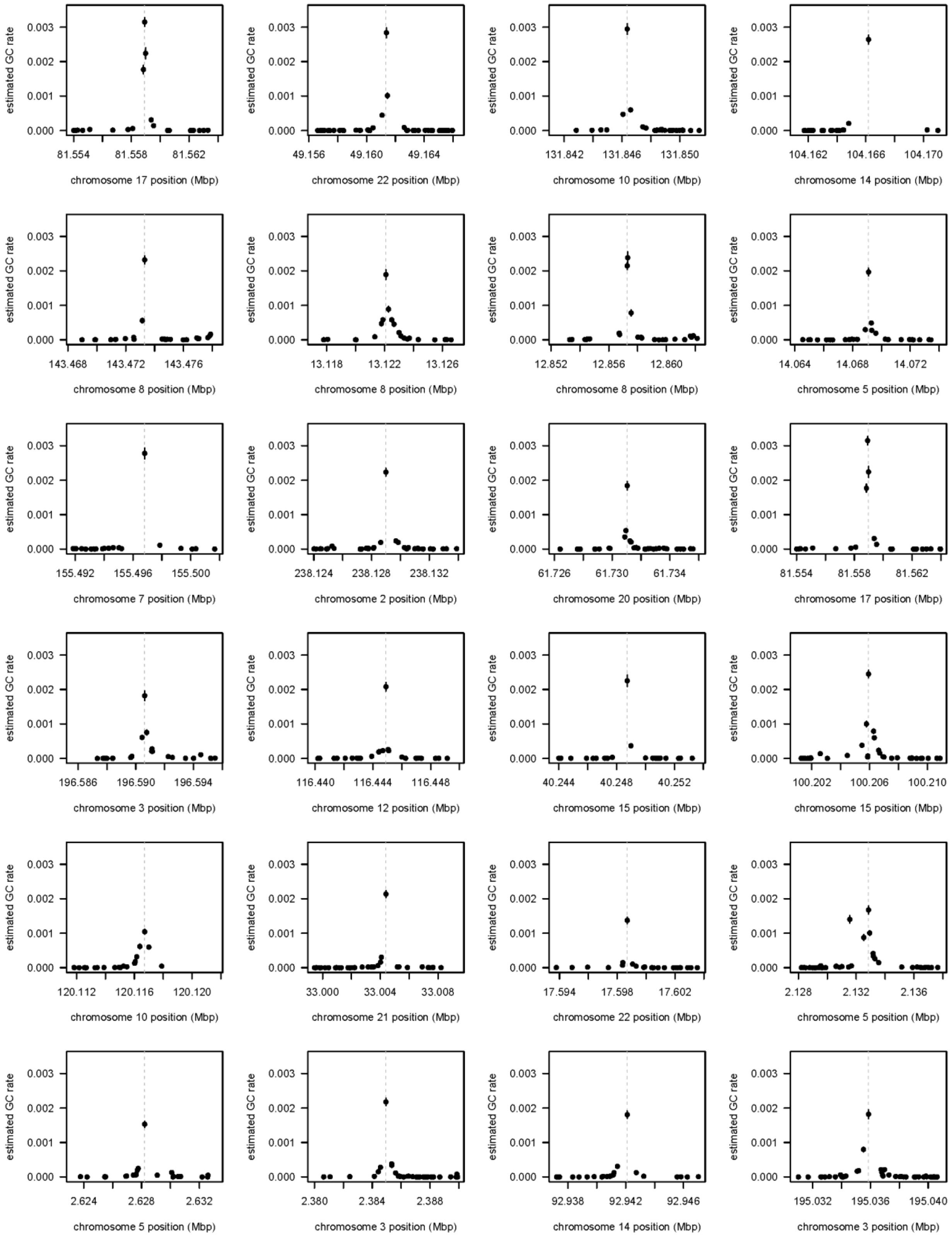
estimated GC rate
0.000
0.001
0.002
0.003
81.554 81.558 81.562
chromosome 17 position (Mbp)
49.156 49.160 49.164
chromosome 22 position (Mbp)
131.842 131.846 131.850
chromosome 10 position (Mbp)
104.162 104.166 104.170
chromosome 14 position (Mbp)
143.468 143.472 143.476
chromosome 8 position (Mbp)
13.118 13.122 13.126
chromosome 8 position (Mbp)
12.852 12.856 12.860
chromosome 8 position (Mbp)
14.064 14.068 14.072
chromosome 5 position (Mbp)
155.492 155.496 155.500
chromosome 7 position (Mbp)
238.124 238.128 238.132
chromosome 2 position (Mbp)
61.726 61.730 61.734
chromosome 20 position (Mbp)
81.554 81.558 81.562
chromosome 17 position (Mbp)
196.586 196.590 196.594
chromosome 3 position (Mbp)
116.440 116.444 116.448
chromosome 12 position (Mbp)
40.244 40.248 40.252
chromosome 15 position (Mbp)
100.202 100.206 100.210
chromosome 15 position (Mbp)
120.112 120.116 120.120
chromosome 10 position (Mbp)
33.000 33.004 33.008
chromosome 21 position (Mbp)
17.594 17.598 17.602
chromosome 22 position (Mbp)
2.128 2.132 2.136
chromosome 5 position (Mbp)
2.624 2.628 2.632
chromosome 5 position (Mbp)
2.380 2.384 2.388
chromosome 3 position (Mbp)
92.938 92.942 92.946
chromosome 14 position (Mbp)
195.032 195.036 195.040
chromosome 3 position (Mbp)

**Figure S2: Gene conversion hotspots.** We selected the markers with the highest estimated gene conversion rates in the TOPMed data and plotted the gene conversion rates in the UK Biobank data at nearby markers. The gray dashed vertical lines give the locations of the hotspots in the TOPMed data. TOPMed hotspots that are within 10 kb of a marker with a larger gene conversion rate are omitted, as are hotspots at markers for which the TOPMed IBD rate is more than 1.4 times or less than 0.6 times the median. TOPMed hotspots for which there is no UK Biobank marker with MAF > 10% within 100 bp or for which there are fewer than 20 UK Biobank markers with MAF > 10% within the 10 kb region centered on the TOPMed hotspot position are also omitted because the plots do not show sufficient detail. Plots are shown in order of TOPMed hotspot rate with highest first, left to right, then top to bottom. The estimated gene conversion (GC) rate at a marker (in the TOPMed data to select the hotspots, and in the UK Biobank data for the y-axis values in these plots) is the number of detected allele conversions divided by the expected heterozygosity of the marker, normalized so that the autosome-wide average is $6 \times 10^{-6}$. Estimates are plotted as dots, while 95% confidence intervals are given as vertical lines through the dots and are obtained by assuming that the number of detected allele conversions follows a Poisson distribution to obtain the standard error and then adding two standard errors to each side of the estimate. Each plot shows all UK Biobank markers with MAF > 10% within 5 kb on either side of the hotspot location. Positions on the x-axes are in GRCh38 coordinates. This figure shows the top 24 hotspots meeting the UK Biobank marker density criteria.

### 4 Supplemental Tables

| $L$ (cM) | $T$ (cM) | IBD rate | IBD FDR | ACs detected | ACs FDR |
|---|---|---|---|---|---|
| 1 | 0.25 | 6.00e-04 | 2.80e-01 | 89793 | 0.045 |
| 1 | 0.5 | 1.10e-04 | 2.30e-02 | 43352 | 0.026 |
| 1.5 | 0.25 | 1.70e-04 | 7.70e-02 | 50038 | 0.027 |
| 1.5 | 0.5 | 7.10e-05 | 9.80e-03 | 30397 | 0.02 |
| 1.5 | 0.75 | 2.30e-05 | 2.70e-03 | 10899 | 0.017 |
| 2 | 0.25 | 4.40e-05 | 2.00e-02 | 18122 | 0.023 |
| 2 | 0.5 | 2.90e-05 | 3.70e-03 | 12109 | 0.018 |
| 2 | 0.75 | 1.70e-05 | 1.20e-03 | 6472 | 0.016 |
| 2 | 1 | 8.60e-06 | 1.90e-04 | 1856 | 0.013 |
| 2.5 | 0.25 | 1.70e-05 | 5.00e-03 | 4576 | 0.020 |
| 2.5 | 0.5 | 1.30e-05 | 1.20e-03 | 3276 | 0.017 |
| 2.5 | 0.75 | 1.00e-05 | 4.40e-04 | 2110 | 0.014 |
| 2.5 | 1 | 7.10e-06 | 1.10e-04 | 1024 | 0.013 |
| 2.5 | 1.25 | 4.60e-06 | 3.90e-05 | 305 | 0.016 |
| 3 | 0.25 | 9.00e-06 | 1.50e-03 | 987 | 0.017 |
| 3 | 0.5 | 7.80e-06 | 3.90e-04 | 761 | 0.018 |
| 3 | 0.75 | 6.60e-06 | 1.20e-04 | 523 | 0.013 |
| 3 | 1 | 5.40e-06 | 2.90e-05 | 325 | 0.009 |
| 3 | 1.25 | 4.20e-06 | 1.60e-05 | 187 | 0.011 |
| 3 | 1.5 | 3.00e-06 | 5.70e-06 | 61 | 0.000 |

**Table S1: Detection rates and false discovery rates for IBD and allele conversions in simulated data.** The simulated data comprise twenty replicates of 10 Mb of sequence data for 10,000 individuals. $L$ and $T$ are the length threshold and trim parameters (in cM) for the multi-individual IBD inference. The IBD rate is the proportion of pairs of haplotypes that are estimated to be IBD averaged across markers. False discovery rate (FDR) for IBD is the proportion of IBD pairs (over all markers) that are not truly pairwise IBD. The number of estimated allele conversions (ACs) is given for each data set. The FDR for ACs is the proportion of estimated ACs that are not true ACs.

| $L$ (cM) | $T$ (cM) | Estimated Ratio | SE |
|---|---|---|---|
| 1 | 0.25 | 1.449 | 0.005 |
| 1 | 0.5 | 1.478 | 0.005 |
| 1.5 | 0.25 | 1.467 | 0.005 |
| 1.5 | 0.5 | 1.480 | 0.006 |
| 1.5 | 0.75 | 1.484 | 0.007 |
| 2 | 0.25 | 1.478 | 0.005 |
| 2 | 0.5 | 1.480 | 0.006 |
| 2 | 0.75 | 1.483 | 0.007 |
| 2 | 1 | 1.483 | 0.008 |

**Table S2: Estimated relative gene conversion rate for simulations with 1.5 times the baseline rate compared to simulations with the baseline rate.** The first column gives the IBD length threshold parameter, the second gives the IBD trim parameter, the third gives the estimated ratio of the estimated gene conversion rates (the true ratio is 1.5), the fourth gives the standard error. The first and last 2 cM of each simulated region are removed before computing the estimated ratios.

| L cM | T cM | TOPMed chr20 10kb | TOPMed chr20 1Mb | TOPMed all 10kb | TOPMed all 1Mb | UKBiobank chr20 10kb | UK Biobank chr20 1Mb |
|---|---|---|---|---|---|---|---|
| 1 | 0.25 | 0.54 | 0.71 | | | | |
| 1 | 0.5 | **0.59** | **0.76** | **0.53** | **0.65** | 0.51 | 0.74 |
| 1.5 | 0.25 | 0.55 | 0.67 | | | | |
| 1.5 | 0.5 | 0.56 | 0.67 | 0.49 | 0.58 | **0.52** | **0.75** |
| 1.5 | 0.75 | 0.53 | 0.58 | 0.43 | 0.54 | | |
| 2 | 0.5 | 0.52 | 0.54 | | | | |
| 2 | 0.75 | 0.50 | 0.44 | | | | |
| 2 | 1 | 0.45 | 0.37 | | | 0.41 | 0.48 |
| 2.5 | 0.75 | 0.45 | 0.35 | | | | |
| 2.5 | 1 | 0.42 | 0.33 | | | | |
| 2.5 | 1.25 | 0.37 | 0.19 | | | 0.32 | 0.44 |

**Table S3: Pearson correlation coefficient between the estimated gene conversion map and the deCODE sex-averaged crossover map.** Results are for TOPMed or UK Biobank data, on chromosome 20 or all autosomes ("all" in column header). Window sizes are 10 kb or 1 Mb and are given in the column header. Initial results were obtained for TOPMed chromosome 20, and a selection of parameter settings were then applied to all autosomes for TOPMed and to chromosome 20 for UK Biobank. Empty cells in the table represent settings not analyzed. The highest correlation in each column is bolded.

| | | 10 kb | | 1 Mb | |
|---|---|---|---|---|---|
| TOPMed $L$ | TOPMed $T$ | UK Biobank $L = 1$ $T = 0.5$ | UK Biobank $L = 1.5$ $T = 0.5$ | UK Biobank $L = 1$ $T = 0.5$ | UK Biobank $L = 1.5$ $T = 0.5$ |
| 1 | 0.25 | 0.834 | 0.823 | 0.957 | **0.958** |
| 1 | 0.5 | **0.860** | 0.856 | 0.951 | 0.949 |
| 1.5 | 0.25 | 0.857 | 0.850 | 0.940 | 0.942 |
| 1.5 | 0.5 | 0.854 | 0.851 | 0.927 | 0.930 |
| 1.5 | 0.75 | 0.848 | 0.845 | 0.885 | 0.891 |
| 2 | 0.5 | 0.847 | 0.843 | 0.863 | 0.869 |
| 2 | 0.75 | 0.837 | 0.834 | 0.857 | 0.862 |

**Table S4. Pearson correlation coefficients between gene conversion rate maps.** Gene conversion rate maps were estimated on chromosome 20 in the TOPMed and UK Biobank data sets. Results are shown for windows of size 10 kb or 1 Mb. The highest correlation for each window size is bolded.

| Window size | Limit on ratio | Number of windows | TOPMed correlation | UK Biobank correlation |
|---|---|---|---|---|
| 10 kb | None | 185083 | 0.56 | 0.43 |
| 10 kb | 10 | 149957 | 0.58 | 0.56 |
| 10 kb | 2 | 96067 | 0.59 | 0.58 |
| 1 Mb | None | 2073 | 0.67 | 0.63 |
| 1 Mb | 2 | 2003 | 0.67 | 0.65 |
| 1 Mb | 1.2 | 1178 | 0.68 | 0.66 |

**Table S5. Correlation between the gene conversion rate maps and the deCODE sex-averaged crossover map.** Data across the autosomes were used, subject to exclusions of windows with low heterozygosity or high or low IBD rate as specified in the main text. If the limit on the ratio is $x$, then windows where the ratio of the TOPMed gene conversion rate to the UK Biobank gene conversion rate is greater than $x$ or less than $1/x$ are ignored. The correlations in the last two columns are between the specified gene conversion rate map and the deCODE 2019 sex-averaged crossover map.

### 6 Supplemental Acknowledgments

We gratefully acknowledge the studies and participants who provided biological samples and data for TOPMed. Funding for the Barbados Asthma Genetics Study was provided by National Institutes of Health (NIH) R01HL104608, R01HL087699, and HL104608 S1. The Mount Sinai BioMe Biobank has been supported by The Andrea and Charles Bronfman Philanthropies and in part by funds from the NHLBI and the National Human Genome Research Institute (NHGRI) (U01HG00638001; U01HG007417; X01HL134588); genome sequencing was funded by contract HHSN268201600037I. The Cleveland Clinic Atrial Fibrillation study was supported by NIH grants R01 HL 090620 and R01 HL 111314, the NIH National Center for Research Resources for Case Western Reserve University and Cleveland Clinic Clinical and Translational Science Award UL1-RR024989, the Cleveland Clinic Department of Cardiovascular Medicine philanthropy research funds, and the Tomsich Atrial Fibrillation Research Fund; genome sequencing was supported by R01HL092577. The Framingham Heart Study was supported by contracts NO1-HC-25195, HHSN268201500001I and 75N92019D00031 from the NHLBI and grant supplement R01 HL092577-06S1; genome sequencing was funded by HHSN268201600034I and U54HG003067. The Hypertension Genetic Epidemiology Network Study is part of the NHLBI Family Blood Pressure Program; collection of the data represented here was supported by grants U01 HL054472, U01 HL054473, U01 HL054495, and U01 HL054509; genome sequencing was funded by R01HL055673. The Jackson Heart Study is supported and conducted in collaboration with Jackson State University (HHSN268201300049C and HHSN268201300050C), Tougaloo College (HHSN268201300048C), and the University of Mississippi Medical Center (HHSN268201300046C and HHSN268201300047C) contracts from NHLBI and the National Institute for Minority Health and Health Disparities (NIMHD); genome sequencing was funded by HHSN268201100037C. The My Life, Our Future samples and data are made possible through the partnership of Bloodworks Northwest, the American Thrombosis and Hemostasis

Network, the National Hemophilia Foundation, and Bioverativ; genome sequencing was funded by HHSN268201600033I and HHSN268201500016C. The Venous Thromboembolism project was funded in part by grants from the NIH, NHLBI (HL66216 and HL83141) and the NHGRI (HG04735). The Vanderbilt Genetic Basis of Atrial Fibrillation study was supported by grants from the American Heart Association (EIA 0940116N), and grants from the National Institutes of Health (HL092217, U19 HL65962, and UL1 RR024975), and by CTSA award (UL1TR000445) from the National Center for Advancing Translational Sciences; genome sequencing was funded by R01HL092577. The Women's Health Initiative program is funded by NHLBI through contracts 75N92021D00001, 75N92021D00002, 75N92021D00003, 75N92021D00004, 75N92021D00005; genome sequencing was funded by HHSN268201500014C.
